# Centenarian Controls Increase Variant Effect-sizes by an average two-fold in an Extreme Case-Extreme Control Analysis of Alzheimer’s Disease

**DOI:** 10.1101/298018

**Authors:** Niccolò Tesi, Sven J. van der Lee, Marc Hulsman, Iris E. Jansen, Najada Stringa, Natasja van Schoor, Hanne Meijers-Heijboer, Martijn Huisman, Philip Scheltens, Marcel J.T. Reinders, Wiesje M. van der Flier, Henne Holstege

## Abstract

The detection of genetic loci associated with Alzheimer’s disease (AD) requires large numbers of cases and controls because variant effect-sizes are mostly small. We hypothesized that variant effect-sizes should increase when individuals who represent the extreme ends of a disease spectrum are considered, as their genomes are assumed to be maximally enriched or depleted with disease-associated genetic variants.

We used 1,073 extensively phenotyped AD cases with relatively young age at onset as extreme cases (66.3±7.9 years), 1,664 age-matched controls (66.0±6.5 years) and 255 cognitively healthy centenarians as extreme controls (101.4±1.3 years). We estimated the effect-size of 29 variants that were previously associated with AD in genome-wide association studies.

Comparing extreme AD-cases with centenarian-controls increased the variant effect-size relative to published effect-sizes by on average 1.90-fold (*SE*=0.29, *p*=9.0×10^−4^). The effect-size increase was largest for the rare high-impact *TREM2 (R74H)* variant (6.5-fold), and significant for variants in/near *ECHDC3* (4.6-fold), *SLC24A4-RIN3* (4.5-fold), *NME8* (3.8-fold), *PLCG2* (3.3-fold), *APOE-ε2* (2.2-fold) and *APOE*-*ε4* (2.0-fold). Comparing extreme phenotypes enabled us to replicate the AD association for 10 variants (*p*<0.05) in relatively small samples. The increase in effect-sizes depended mainly on using centenarians as extreme controls: the average variant effect-size was not increased in a comparison of extreme AD cases and age-matched controls (0.94-fold, *p*=6.8×10^−1^), suggesting that on average the tested genetic variants did not explain the extremity of the AD-cases. Concluding, using centenarians as extreme controls in AD case-controls studies boosts the variant effect-size by on average two-fold, allowing the replication of disease-association in relatively small samples.

## Introduction

Alzheimer’s disease (AD) is characterized by a slow but progressive loss of cognitive functions, leading to loss of autonomy.^1^ AD is rare at the age of 65 years, but its incidence increases exponentially to 40% at the age of 100 years.^2^ It is currently the most prevalent cause of death at old age and one of the major health threats of the 21st century.^1^ Better understanding of the etiological factors that determine AD is warranted as no treatment is currently available. Heritability plays an important role as genetic factors are estimated to determine 60-80% of the risk of AD.^3^ About 30% of the genetic risk is attributable to the *ε4* allele of *APOE* gene, and large collaborative efforts have identified over two dozen additional genetic loci that are associated with a slight modification of the risk of AD.^4–17^ The design of these association studies relies on the comparison of very large numbers of cases with age-matched controls, such that detected associations can be attributed specifically to the disease.^18^ However, given the prevalence of AD in the aging population, it is likely that a significant fraction of the controls will develop the disease at a later age. Therefore, as the AD risk for future cases likely involves the same genetic variants, using age-matched controls may quench variant association signals. This may, in part, explain the mostly small variant effect-sizes associated with common variants. Also, GWAS studies mostly compare common genetic variants that are widely propagated in the population; as a consequence, these have mostly small effects on AD risk.^19^ Rare genetic variants often have larger effect-sizes than common variants, but as there are fewer carriers available in the population, the requirement for large sample sizes stands.^20^

Instead of increasing sample sizes of genetic studies to detect novel disease-associated genetic loci, an alternative strategy is to increase variant effect-sizes by sampling individuals with extreme phenotypes.^20–22^ For AD and other age-related diseases, extreme cases may be defined by having a relatively early age at disease-onset, and having the phenotypic features characteristic for the disease, as defined by diagnostic assessment. Extreme controls are represented by individuals who reach extreme ages without the disease.^21,23,24^ Indeed, in a case-control study of type-2 diabetes, the effect-sizes for variants that were previously associated with the disease were increased when using centenarians as extreme controls.^23^ The effect of using extreme phenotypes in other age-related diseases has not been studied.

Here, we explored the potential of using extreme phenotypes for genetic studies of Alzheimer’s disease (AD) by investigating the change in effect-size of known AD-associated variants. Furthermore, using an age- and population-matched reference group, we investigated the contribution of each extreme phenotype.

## Methods

### Cohort description

As extreme AD cases group (denoted by *EA*) we used 1,149 AD cases from the Amsterdam Dementia Cohort (ADC). The ADC comprises patients who visit the memory clinic of the VU University Medical Center, The Netherlands.^25,26^ This cohort of AD patients is extensively characterized and comprises 503 early-onset cases (denoted by *eEA*) with an age at onset <65 years, and 646 late-onset cases (denoted by *lEA*). At baseline, all subjects underwent a diagnostic assessment including neurological examination, standard laboratory tests of blood and cerebrospinal fluid, electroencephalogram and brain magnetic resonance imaging. Clinical diagnosis is made in consensus-based, multidisciplinary meetings. The diagnosis of probable AD was based on the clinical criteria formulated by the National Institute of Neurological and Communicative Disorders and Stroke - Alzheimer’s Disease and Related Disorders Association (NINCDS-ADRDA) and based on National Institute of Aging - Alzheimer association (NIA-AA).^25–28^

As extreme control group (denoted by *EC*) we used 268 self-reported cognitively healthy centenarians from the 100-plus Study cohort.^29^ This study includes Dutch-speaking individuals who (i) can provide official evidence for being aged 100 years or older, (ii) self-report to be cognitively healthy, which is confirmed by a proxy, (iii) consent to donation of a blood sample, (iv) consent to (at least) two home visits from a researcher, and (v) consent to undergo an interview and neuropsychological test battery.

As ‘normal controls’ (denoted by *NC*) we used 1,717 middle-aged (55-85 year-old) individuals from a representative sample of Dutch individuals from the Longitudinal Aging Study Amsterdam (LASA) cohort.^30,31^ LASA is an ongoing longitudinal study of older adults initiated in 1991, with the main objective to determine predictors and consequences of aging.

The Medical Ethics Committee of the VU University Medical Center (METC) approved the ADC cohort, the LASA study and the 100-plus Study. All participants and/or their legal guardians gave written informed consent for participation in clinical and genetic studies.

### Genotyping and imputation of 29 selected AD-associated genetic variants

We selected 29 single nucleotide variants for which evidence for a genome-wide significant association with Alzheimer’s disease was found in previous studies (*Table S1*).^4–17^ Genetic variants were determined by standard genotyping or imputation methods. Briefly, we genotyped all individuals using the Illumina Global Screening Array (GSAsharedCUSTOM_20018389_A2) and applied established quality control methods.^32^ We used high quality genotyping in all individuals (individual call rate >98%, variant call rate >98%), individuals with sex mismatches were excluded and HWE-departure (d-HWE) was considered significant at *p*<1×10^−6^. Genotypes were prepared for imputation using provided scripts (HRC-1000G-check-bim.pl).^33^ This script compares variant ID, strand and allele frequencies to the haplotype reference panel (HRC v1.1, April 2016).^33^ Finally, all autosomal variants were submitted to the Michigan imputation server (https://imputationserver.sph.umich.edu).^32^ The server uses *SHAPEIT2* (v2.r790) to phase data and imputation to the reference panel (v1.1) was performed with *Minimac3*.^32,34^ A total of 1,149 extreme AD cases, 1,717 normal controls and 268 extreme (centenarian) controls passed quality control. Prior to analysis, we excluded individuals of non-European ancestry (*N_EA_* = 67, based on 1000Genomes^35^ clustering) and individuals with a family relation (*N_EA_* = 9, *N_EC_* = 13, *N_NC_* = 53, identity-by-descent ≥ 0.3),^36^ leaving 1,073 extreme AD cases (*N_eEA_* = 464 and *N_lEA_* = 609), 1,664 normal controls and 255 centenarian controls for the analysis.

### Statistical Analysis

For each AD-associated variant, we explored the *change in effect-size* (*E*) relative to reported effect-sizes when 1) comparing extreme AD cases with extreme (centenarian) controls (*EA vs*. *EC*); 2) comparing extreme AD cases with normal controls (*EA vs*. *NC*); and 3) comparing normal AD cases with extreme (centenarian) controls (*NA vs. EC*). To calculate variant effect-sizes, we used logistic regression models correcting for population stratification (principal components 1 to 6).^37,38^ We calculated odds ratios (OR) relative to the HRC alternative allele assuming additive genetic effects, and estimated 95% confidence intervals (CIs).

We estimated the *change in effect-size* relative to reported effect sizes (*E*) as follows:

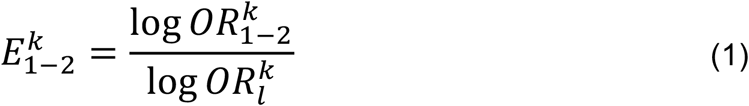

Where 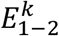 indicates the effect-size change for variant *k* in a comparison of cohort 1 and cohort 2, *e.g*, *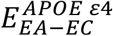* indicates the effect-size change for the *APOE ε4* variant when extreme AD cases (*EA*) are compared with cognitive healthy centenarians (*EC*). The log 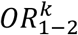 denotes the *effect-size* of variant *k* when comparing cohort 1 and cohort 2. The effect-size of variant *k* reported in literature (*Table S1*) is denoted by log 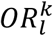.

We estimated the added value of using extreme (centenarian) controls rather than ‘normal age-matched controls’ in a case-control analysis. For this, we wanted to compute the change in effect size when comparing non-extreme AD cases with extreme controls (*NA* vs *EC*). As we do not have direct access to ‘normal AD cases’, we estimated the effect-size for the *NA-EC* comparison by summing (1) the effect-size when comparing ‘normal AD cases’ and ‘normal controls’, as reported in literature (log 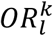), and (2) the effect size when comparing normal controls (*NC*) with extreme (centenarian) controls (*NC* vs *EC*), *i.e*. 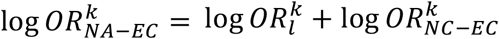. The added value of using extreme controls in a case control analysis then becomes:

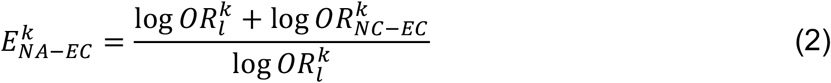

To assess whether age at disease onset had an impact on the change in effect-size due to the extreme cases (*E_EA-NC_*), we estimated the log 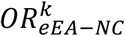 (early-onset extreme AD cases *vs*. normal controls) and log 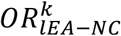 (late-onset extreme AD cases *vs*. normal controls), and their 95% confidence intervals. Then, we computed the probability that the effect size changes 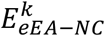 and 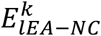 differed using a two-samples z-test (two-tailed *p-value*).

### Determining significance of change in effect-size

For each variant, we estimated 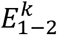 and a 95% confidence intervals (CI) by sampling (*S*=10,000) from the log 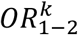 and log 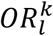 based on their respective standard errors. The probability of divergence between the distributions of the log 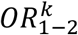 and the log 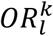 was determined using a two-sample z-test (two-tailed *p-value*).

The probability of observing 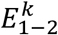 > 1, *i.e* an increased effect-size for variant *k*, is considered to be a Bernoulli variable with *p*=0.5 (equal chance of having an increased/decreased effect). The number of variants that show an increase in effect (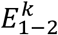 > 1) then follows a binomial distribution

The average change in effect-size across all *K*=29 tested variants is calculated as follows:

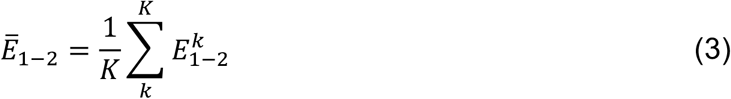

Confidence intervals and probability of divergence between 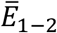 and previously reported effect-sizes were estimated by sampling (*S*=10,000, two-tailed *p-value*).

All described statistical analysis were performed with *PLINK* (v1.90b4.6) or *R* (v3.3.2).^39,40^

## Results

After quality control of the genetic data, we included 1,073 extreme AD cases (with mean age at onset 66.4±7.8 and 52.7% females), 1,664 normal (age-matched) controls (mean age at inclusion 66.0±6.5, 53.7% females), and 255 cognitive healthy centenarians as extreme controls (mean age at inclusion 101.4±1.3, 74.7% females) (*Table 1*). Within the extreme AD cases group, there were 464 early-onset cases (mean age at onset 59.1±4.1, 54% females), and 609 late-onset cases (mean age at onset 72.1±4.8, 51% females). The age at onset of the extreme AD cases was on average 8.2 years earlier compared to previous GWA studies; the age at disease onset was on average 15.4 years earlier in early-onset cases and 2.5 years earlier in late-onset cases, while the age of study inclusion of our centenarian controls was on average 29.5 years higher than for previously published controls (*Figure 1*).

**Table 1:** Population characteristics.

|  | **Extreme AD**<br>**Cases (EA)** | **Centenarian**<br>**controls (EC)** | **Normal controls**<br>**(NC)** |
| --- | --- | --- | --- |
| **Number of individuals** | 1,073 | 255 | 1,664 |
| **Females (%)** | 564 (52.6) | 191 (74.9) | 893 (53.7) |
| **Age (SD)\*** | 66.4 (7.8) | 101.4 (1.3) | 66.0 (6.5) |
| **ApoE $\epsilon 4$ (%)** | 981 (42.7) | 44 (8.6) | 533 (16.0) |
| **ApoE $\epsilon 2$ (%)** | 76 (3.5) | 78 (15.3) | 304 (9.1) |

\*Age at onset for extreme Alzheimer's disease cases, age at study inclusion for extreme controls and normal controls; Abbreviations: SD, standard deviation; ApoE, Apolipoprotein E allele count for $\epsilon 4$ and $\epsilon 2$ , respectively; Reference to the cohorts reported in this table are: <sup>25,26,29,30</sup>

**Figure 1:**
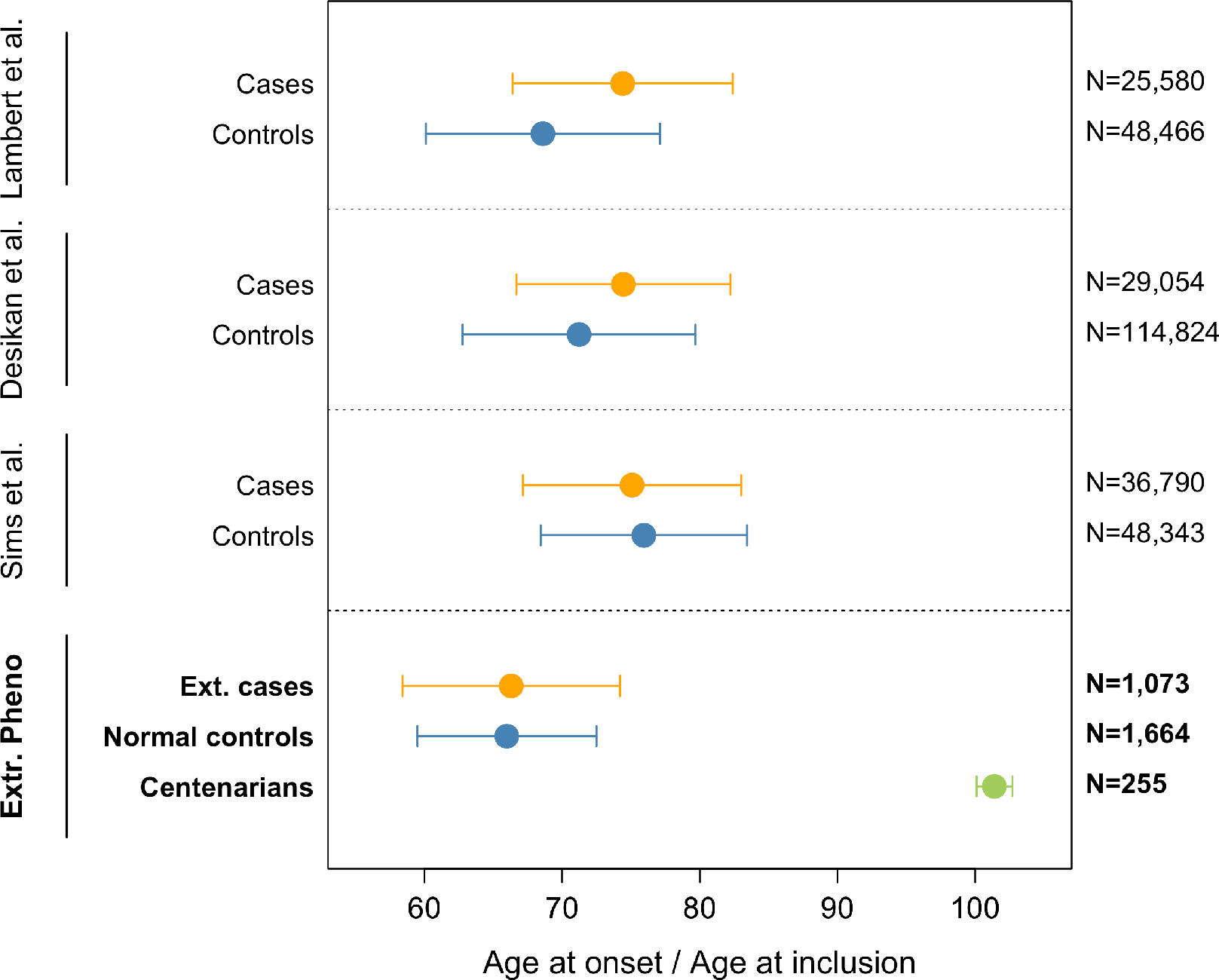
Comparison of age at disease-onset and age at inclusion for cases and controls in previously reported case-control comparisons, and in our extreme phenotypes comparison. Weighted mean and (combined) standard deviation of the age at onset for AD cases and age at inclusion for controls. As weights, we used the sample sizes of each GWA study. Note that previous case-control studies of AD included samples from multiple cohorts, sometimes overlapping across studies. References to the cohorts reported in this figure are: ^7,8,13,25,26,30^

### Effect of comparing extreme cases and centenarian controls

In a genetic comparison of extreme AD cases and centenarian controls (*EA-EC* comparison) the average effect-size over all 29 genetic variants was 1.90-fold increased relative to the effect-sizes reported in published studies (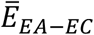 = 1.90±0.29; *p* = 9.0×10^−4^) (*Figure 3*). For 21 out of 29 variants, we observed an increased effect size (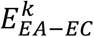 > 1), which is significantly more than expected by chance (*p* = 1.2×10^−2^) (*Figure 2* and *Table 2)*. The increase in effect-size ranged from 1.06 (variant near *CASS4*) to 6.46 (variant in *TREM2 [R47H]*). For variants near or in the genes *TREM2 (R47H), SLC24A4-RIN3* and *ECHDC3*, the increase was more than 4-fold compared to previously reported effect-sizes. For 9 variants the effect-size increase was 2-4-fold (in or near the genes *NME8*, *PLGC, HLA-DRB1, 2CD2AP, ZCWPW1, ABCA7 APOE [ε2], [A>G], HS3ST1* and *ABI3*, in order from high to low effect-size increases). For 9 variants the increase was between 1 and 2-fold (in or near genes, *APOE ε4, RPHA1, CELF1, PTK2B, MS4A6A, SORL1, BIN1, PICALM* and *CASS4*) (*Figure 2*). The effect-sizes for 6 genetic variants were not increased in our extreme phenotype analysis compared to previously reported effect-sizes (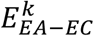 between 0 and 1): in or near *TREM2 [R62H], KANSL1 CR1, ABCA7 [G>C], CLU*, and *INPP5D*). Lastly, the effect-sizes of 2 variants were in the opposite direction compared to previously reported effects (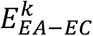 > 0). Specifically, for the variant in *FERMT2* we found an inverted direction of effect-size and a lower magnitude of effect as compared with previous studies (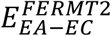 between 0 and −1). For the variant near *MEF2C* we observed a larger effect-size as compared to those previously published, but in the opposite direction (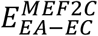> −1).

Overall, for 7 variants, the effect size was significantly increased relatively to the previously reported effect-sizes (*Table 2*), in or near genes *APOE ε2* (2.2-fold, *p* = 1.4×10^−7^), *APOE ε4* (2.0-fold, *p* = 1.5×10^−9^), *SLC24A4-RIN3* (4.5-fold, *p* = 1.6×10-^3^), *ECHDC3* (4.6-fold, *p* = 1.1×10^−2^), *PLCG2* (3.3-fold *p* = 1.4×10^−2^), *NME8* (3.9-fold, *p* = 1.7×10^−2^) and *MEF2C* (−1.9-fold, *p* = 1.8×10^−2^). Variants with significant effect-size changes, were also more likely to be associated with AD in a comparison of extreme cases and centenarians. The association with AD reached nominal significance (*p<*0.05) in 10 out of 21 variants with a changed effect-size (*Table 2*). Next to *APOE ε4* (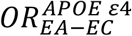 = 2.1, SE = 0.17, *p* = 1.3×10^−33^) and *APOE ε2* (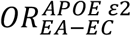 = −1.8, *p* = 3.2×10^−21^), variants in or near these genes were significantly associated with AD: *SCL24A4-RIN3, PLCG2, ECHDC3, NME8, BIN1, ZCWPW1, ABCA7 (A>G)* and *HLA-DRB1* (*Table 2*).

**Figure 2:**
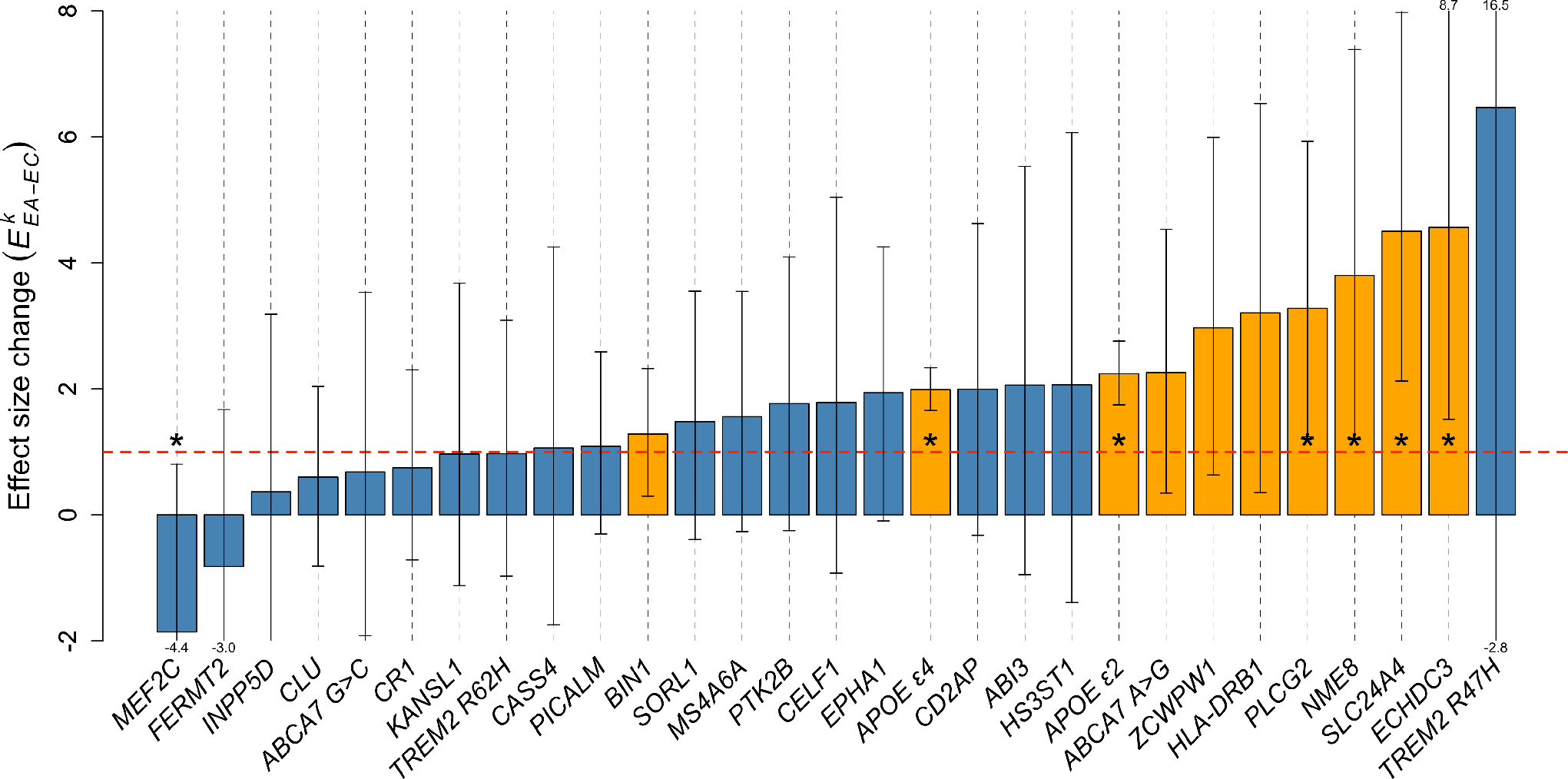
Change in variant effect-size using extreme cases and centenarian controls relative to published effect-sizes, for 29 AD associated genetic variants. Dashed red line at 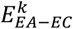=1 indicates same effect-size as reported in literature. Orange bars indicate nominal statistical significance for the association with AD (*p*<0.05). Stars indicate significant changes of effect-size relative to previously reported effect-sizes (*p*<0.05, two-sample z-test).

**Table 2:**
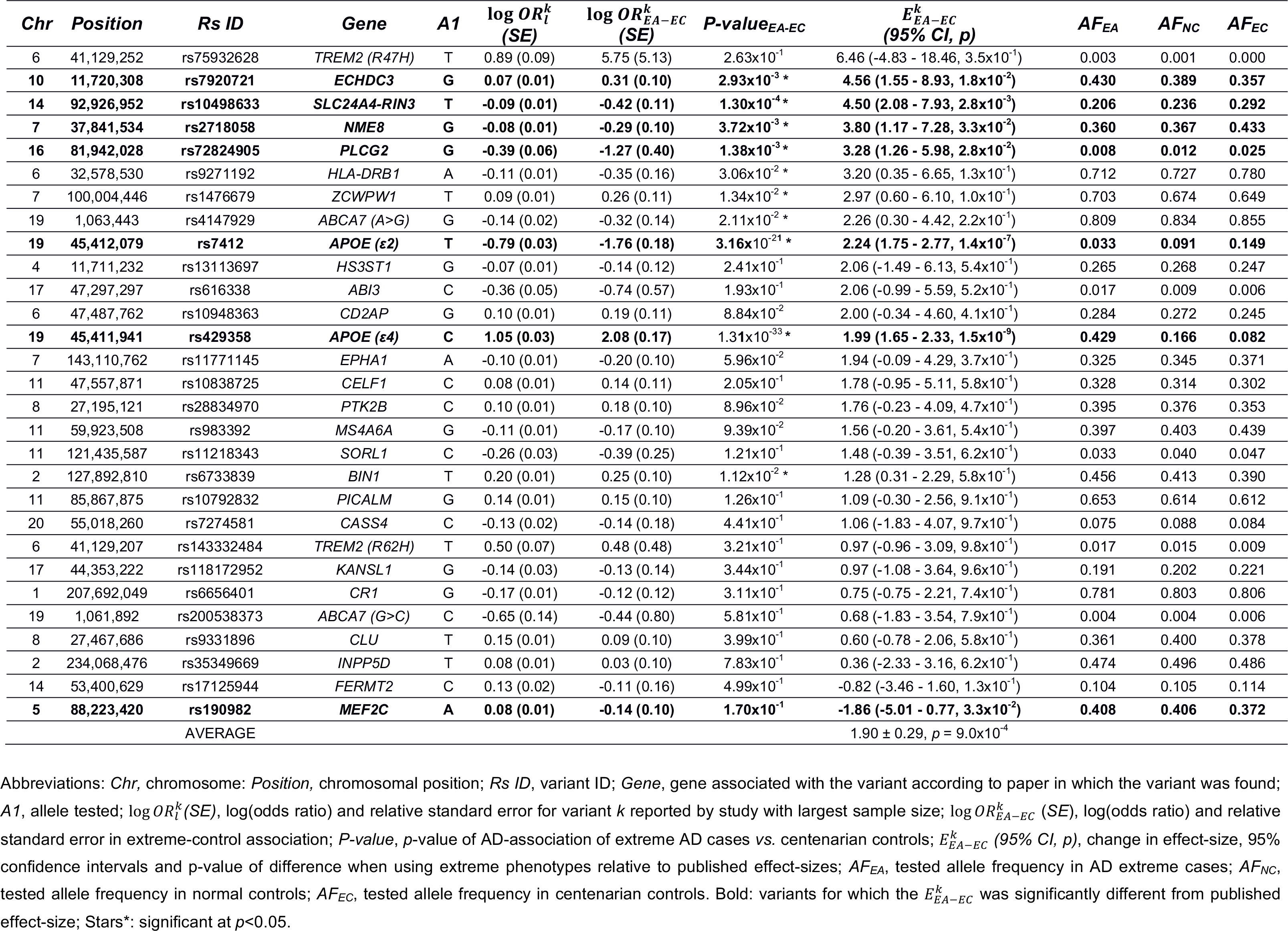
Association statistics of the 29 tested AD-associated variants.

### Effect of using extreme AD cases

The average effect-size in a comparison of extreme AD cases with normal controls (*EA vs. NC*) did not significantly change relative to the previously reported effect-sizes (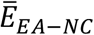 = 0.94±0.12, *p*=6.8×10^−1^) (*Figure 3*). For 14 individual variants, we observed an increased effect size, which was expected by chance (*p*=0.5), *Figure S1* and *Table S2*). The effect size was significantly increased for *APOE ε4* variant (1.3-fold, *p* = 1.4×10^−5^), and nominally significant for *APOE-ε2* (1.4-fold, *p* = 1.7×10^−2^).

**Figure 3:**
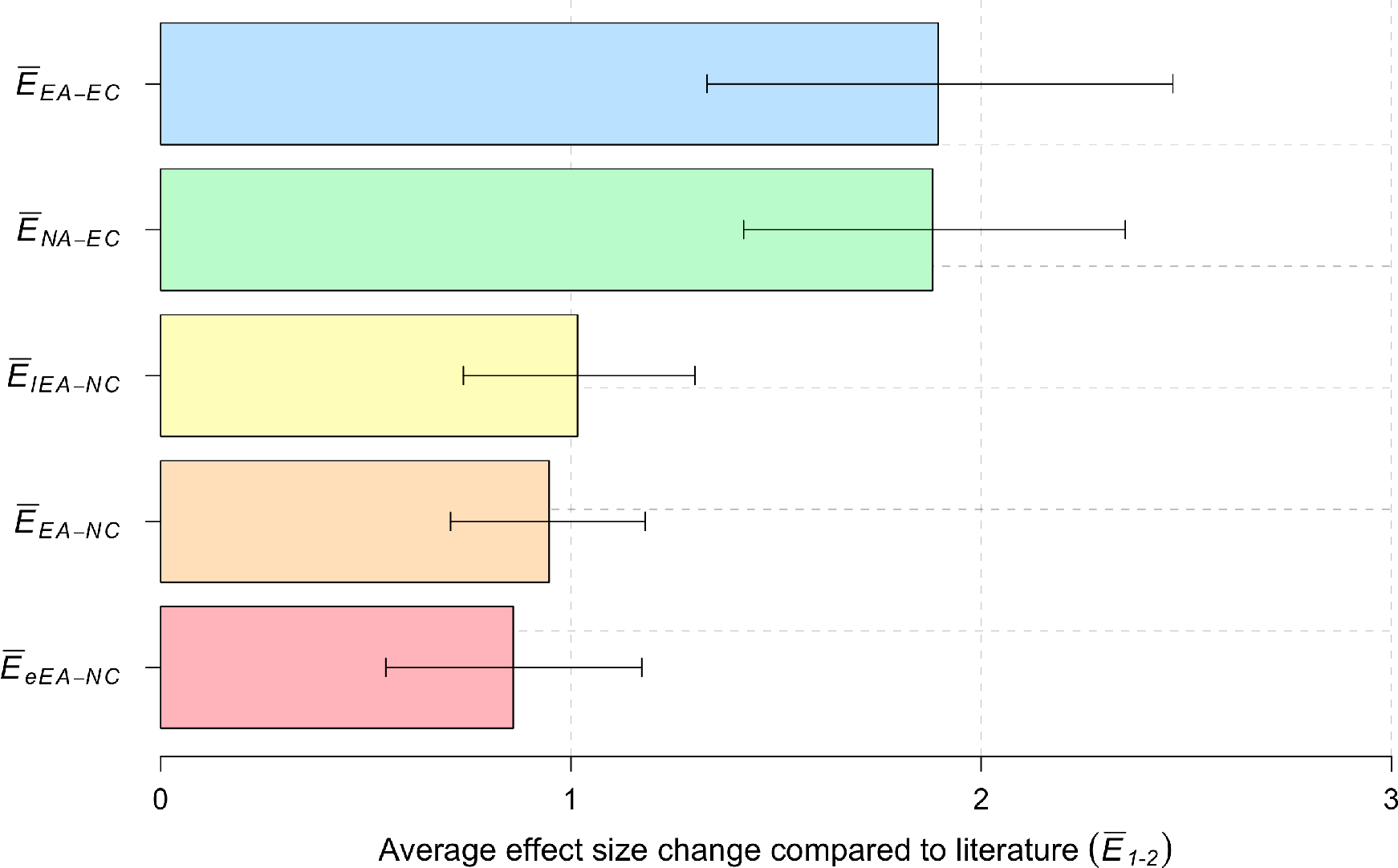
Average increase in effect-size for the different comparisons. Average increase in effect sizes for: Extreme AD cases (*N_EA_*=1,073), of which early onset cases (*N_eEA_*=464) late onset cases (*N_lEA_*=609); centenarian controls (*N_EC_*=255); normal controls (*N_NC_*=1,664). 95% confidence intervals were estimated by random sampling (*S=*10,000).

We then separated AD cases into early-onset extreme AD cases (*N_eEA_* = 464) and late-onset extreme AD cases (*N_lEA_* = 609), and estimated the change in effect-sizes. When using early onset cases the average effect-size was lower relative to previously published effect sizes (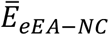 was 0.86±0.16 (*p* = 7.9×10^−1^), while for late-onset cases the effect size was similar to published effect sizes (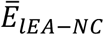 was 1.01±0.14, *p* = 4.6×10^−1^) (*Figure S3* and *Table S3*). We found significant differences between the effect-sizes in early-onset and late-onset AD cases (log 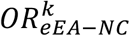 and log 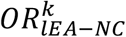, respectively) for the variants in or near *APOE ε2* (−1.41 *vs*. −0.89; *p*=5.0×10^−2^), *ZCWPW1* (0.01 *vs*. 0.24: *p*=1.6×10^−2^) and *MS4A6A* (0.12 *vs*. −0.13; *p*=7.9×10^−3^).

### Effect of extreme controls

In a comparison of normal AD cases and extreme (centenarian) controls (*NA vs*. *EC*), the effect-size was on average 1.88-fold higher relative to previously reported effect-sizes (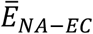 = 1.88±0.24, *p* = 1.0×10^−4^) (*Figure 3*). This was almost identical to the average increase in effect-size when we compared the extreme cases with centenarian controls (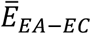 = 1.90±0.29; *p* = 9.0×10^−4^) (*Figure 3*). At the variant level, the change in effect-sizes was similar in both analyses, with the exception of the rare *TREM2 (R47H)* variant, whose effect-size increase was higher in the comparison with the extreme cases (but with high confidence intervals) (*Figure S4-A*).

In further concordance with the comparison of the extreme phenotypes, we observed an increased effect-size for 24/29 variants relative to published variant effect-sizes (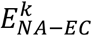 > 1), which is more than expected by chance (*p* = 2.7×10^−4^) (*Figure S2* and *Table S2*). We found a significant increase in effect-size for variants in or near *APOE-ε2* (1.7-fold, *p* < 5×10^−5^), *APOE-ε4* (1.7-fold, *p* < 5×10^−5^), *NME8* (4.5-fold, *p = 3.5×10^−3^*), *SLC24A4-RIN3* (3.9-fold, *p = 4.5×10^−3^*) and *PLCG2* (2.9-fold, *p = 1.9×10^−2^*). In line with this, for almost all variants individually, the extreme controls contributed more to the effect size change in the extremes-comparison, than the extreme cases (*Figure S4-B*).

## Discussion

In this study, we found that the effect sizes of 29 variants previously identified in genetic case-control analyses for Alzheimer’s Disease were increased in a case-control analysis of extreme phenotypes. The use of extreme AD-cases and cognitively healthy centenarians as extreme-controls increased effect sizes for association with AD up to 6-fold, relative to previously published effect-sizes. On average, the use of extreme phenotypes almost doubled the variant effect-size. Although changes in effect-size were different per variant, the effect-size increase was driven mainly by the centenarian controls. This profound increase enabled us to replicate the association with AD of 10 variants in relatively small samples.

In a comparison of AD cases (either normal or extreme) with centenarian controls, we observed significant effect-size increases for variants in or near *PLCG2, NME8, ECHDC3, SLC24A4-RIN3, APOE-ε2* and *APOE-ε4*. This suggests that the tested variants or loci might contribute to the long-term preservation of cognitive health and/or to longevity in general. *PLCG2* and *NME8* are implicated in immunological processes,^8,43^ while *SLC24A4*, *ECHDC3* and *APOE* are involved in lipid and cholesterol metabolism.^17,44,45^ Both these processes were previously associated with longevity,^46,47^ such that an overlapping etiology of maintained cognitive health and maintained overall health may contribute to the observed increase in effect-size. However, with the exception of the *APOE* locus, these loci were thus far not associated with longevity in GWAS studies.^48–51^ We speculate that the association might be dependent on the cognitive health in the centenarians of the 100-plus Study cohort.^29^ Alternatively, longevity studies may have been underpowered to detect the association of these loci with extreme survival. Future studies will have to establish the mechanism behind the association of these genes with preserved cognitive health. Next to *APOE*, the *HLA-DRB1* locus has been associated with both AD^13^ and longevity.^48^ However, its most informative variants, *rs9271192* for AD and *rs34831921* for longevity, are not in linkage disequilibrium,^52^ suggesting that these are independent signals.

Using extreme cases did not increase the variant effect-sizes relative to published effect-sizes, even though most of the extreme cases were biomarker confirmed and their mean age at onset was 8.2 years younger than the mean age at onset in other studies.^7,8,13^ This suggests that based on the tested genetic variants, the “phenotypically extreme” cases presented in this study were not genetically more extreme than cases presented in other studies.

Counter intuitively, in a comparison with normal controls, the variant effect-sizes of early-onset AD cases were on average *lower* than the variant effect-size of late-onset AD cases. One explanation for this observation may be that the early age at onset may have been driven by rare, high-impact variants,^19^ while the disease onset at later ages may depend to a greater extent on more common risk variants, which are tested here. However, at the variant level, we found significant differences between the effect-sizes in early-onset and late-onset cases for variants in/near *ZCWPW1* and *APOE ε2*, and also in —opposite directions— for the variant in *MS4A6A*. These results are a first indication that these variants may differentially influence age of disease onset, however, future experiments will have to confirm this finding.

Here, we find that the majority of the observed increase in effect-size in a genetic case-control study of extreme phenotypes is attributable to the extreme controls. We note that the centenarians used in this study were selected for their preserved cognitive health, which might have further enlarged the effect-size increase for genetic variants that were previously identified for their AD-association. We acknowledge that using centenarians as controls in genetic studies of AD could result in the detection of variants associated with extreme longevity, such that newly detected AD-associations need to be verified in an age-matched AD case-control setting. Nevertheless, the effect-sizes for all but two variants are in the same direction as previously reported, which suggests that the tested AD variants do not have significant pleiotropic activities that counteract their AD-related survival effects. Notably, the two variants with an opposite effect in the comparison of the extremes relative to published effect sizes, in or near *MEF2C* and *FERMT2*, also did not associate with AD in our age-matched case-control analysis. This suggests that the AD-association of the *MEF2C* and *FERMT2* variants might be false positive findings in previous studies. This is in line with results from unpublished GWASs of AD in which AD-associations of variants near the *MEF2C* and *FERMT2* genes were not replicated^41,42^ (*p* = 5.4×10^−3,41^ *p* = 3.0×10^−4^ for *MEF2C*^42^ and *p* = 1.6×10^−5^ for *FERMT2*^42^ variant, with 5.0×10^−8^ being the genome-wide significance threshold). An additional strength of our study is that our cohorts of AD patients and controls, were not previously used in the discovery of any of the known AD associated variants;^4–17^ we thus provide independent replication in a genetically homogeneous group of individuals, as they all came from one specific population (Dutch).

Concluding, in our comparison of cases and controls with extreme phenotypes we found that on average, the effect of AD-related variants in genetic association studies almost doubled, while at the variant level effect-sizes increased up to six-fold. The observed increment in effect-size was largely driven by the centenarians as extreme controls, identifying centenarians as a valuable resource for genetic studies, with possible applications for other age-related diseases.

## Acknowledgements

Research of the VUmc Alzheimer center is part of the neurodegeneration research program of Amsterdam Neuroscience (www.amsterdamresearch.org). The VUmc Alzheimer Center is supported by Stichting Alzheimer Nederland (WE09.2014-03) and Stiching VUmc fonds. The clinical database structure was developed with funding from Stichting Dioraphte (VSM 14 04 14 02). The Dutch case-control study is part of EADB (European Alzheimer DNA biobank) funded by JPcofundNL (ZonMW project number: 733051061). The authors declare no conflict of interests.

